# Transcription factor binding sites are frequently under accelerated evolution in primates

**DOI:** 10.1101/2022.04.29.490094

**Authors:** Xinru Zhang, Yi-Fei Huang

**Affiliations:** Department of Biology, Pennsylvania State University, University Park, PA 16802, USA; Huck Institutes of the Life Sciences, Pennsylvania State University, University Park, PA 16802, USA; Bioinformatics and Genomics Graduate Program, Pennsylvania State University, University Park, PA 16802, USA

## Abstract

Recent comparative genomic studies have identified many human accelerated elements (HARs) with elevated substitution rates in the human lineage. However, it remains unknown to what extent transcription factor binding sites (TFBSs) are under accelerated evolution in humans and other primates. Here, we introduce two pooling-based phylogenetic methods with dramatically enhanced sensitivity to examine accelerated evolution in TFBSs. Using these new methods, we show that more than 6,000 TFBSs annotated in the human genome have experienced accelerated evolution in Hominini, apes, and Old World monkeys. Although these TFBSs individually show relatively weak signals of accelerated evolution, they collectively are more abundant than HARs. Also, we show that accelerated evolution in Pol III binding sites may be driven by lineage-specific positive selection, whereas accelerated evolution in other TFBSs might be driven by nonadaptive evolutionary forces. Finally, the accelerated TFBSs are enriched around neurodevelopmental and pluripotency genes, suggesting that accelerated evolution in TFBSs may drive the divergence of neurodevelopmental processes between primates.

## Introduction

During the course of evolution, a subset of genes and regulatory elements may be subject to different pressures of natural selection in distinct species. These genomic elements often have varying substitution rates across species, which may be identified by phylogenetic models with lineage-specific substitution rates^1–10^. Notably, previous studies have revealed a few thousand human accelerated regions (HARs) with dramatically elevated substitution rates in the human lineage compared to other vertebrates^7–15^. A large proportion of HARs are neural enhancers^14–20^ and frequently subject to strong positive selection in the human lineage^9–11, 21^, suggesting that they may contribute to the adaptive evolution of human brain. Also, recent studies show that deleterious mutations in HARs may be associated with neurodevelopmental disorders^22–26^, highlighting the key role of HARs in maintaining the integrity of the central nervous system. Thus, characterizing genomic elements under accelerated evolution is of great importance for understanding the genomic basis of human evolution and disease.

While numerous studies have been conducted to examine accelerated evolution in humans and other species^7–15, 27–29^, the existing studies may suffer from two critical limitations. First, most of the previous studies have focused on conserved noncoding elements under accelerated evolution. Because a large proportion of noncoding regulatory elements may be subject to frequent evolutionary turnover^30–34^, these studies may not be able to characterize accelerated evolution in non-conserved regulatory elements. Second, the previous studies have focused on identifying individual HARs with a genome-wide level of significance. Because of the small amount of alignment data in a single genomic element and the high burden of multiple testing correction associated with a genome-wide scan, these studies may have limited statistical power to detect weakly accelerated evolution driven by relaxed purifying selection or weak positive selection. Altogether, it remains unknown to what extent non-conserved genomic elements are subject to weakly accelerated evolution.

Substitutions in transcription factor binding sites (TFBSs) are a main driver of phenotype diversity between species^35–37^, implying that TFBSs may also be subject to accelerated evolution. However, to the best of our knowledge, accelerated evolution in TFBSs has not been systematically explored in previous studies, possibly because the majority of TFBSs are not highly conserved across vertebrates^30–33, 38, 39^. Also, TFBSs might be subject to weaker acceleration compared to conserved elements because the phenotypic effects of mutations are weaker in TFBSs than in conserved elements^40^. Therefore, previous phylogenetic methods dedicated to infer strong signals of accelerated evolution may be underpowered to detect TFBSs under weakly accelerated evolution.

Here, we introduce two novel phylogenetic methods for exploring TFBSs under accelerated evolution. Unlike previous methods that analyze individual elements separately^7–15^, our new approaches pool thousands of TFBSs with similar functions together to boost the statistical power to detect weak signals of accelerated evolution. These new methods allow us to rigorously test whether a group of TFBSs as a whole is significantly enriched with accelerated elements, despite that we may lack statistical power to identity individual TFBSs under accelerated evolution due to limited alignment data in a single TFBS. Using these methods, we show that TFBSs of numerous transcription factors are likely to be under accelerated evolution in Hominini, apes, and Old World monkeys. Compared to previously identified HARs, these TF-BSs show weaker acceleration but are more abundant genome-wide. Among these accelerated TFBSs, binding sites of DNA-directed RNA polymerase III (Pol III) show the strongest signal of acceleration, which might be driven by strong lineage-specific positive selection on par with HARs. Taken together, accelerated evolution may be a common characteristic of TFBSs in Hominini, apes, and Old World monkeys.

## Results

### Pooling-based phylogenetic inference of accelerated evolution

In the current study, we introduce a novel software application, GroupAcc, which include two pooling-based phylogenetic approaches with improved statistical power to infer weakly accelerated evolution. The key idea of GroupAcc is to group TFBSs by the bound transcription factor and then examine whether each TFBS group as a whole shows an elevated substitution rate in a lineage of interest. By pooling alignment data from a large number of TFBSs, our new methods have significantly higher statistical power to detect weakly accelerated evolution at the group level even if the signals of acceleration are statistically insignificant at the level of individual TFBSs.

In the first method, we utilize a *group-level likelihood ratio test (LRT)* to infer whether a TFBS group as a whole shows an elevated substitution rate in a predefined foreground lineage compared to the other lineage (background lineage) (Fig. 1). To this end, we first fit a reference phylogenetic model to the concatenated alignment of all TFBSs, where we estimate the branch lengths of a phylogenetic tree, the gamma shape parameter for rate variation among nucleotide sites^41^, and the parameters of the general time reversible substitution model^42^. Assuming that the majority of TFBSs may not be under accelerated evolution, the reference phylogenetic model may represent the overall pattern of sequence evolution in TFBSs when accelerated evolution is absent. Given the reference phylogenetic tree, we then fit the group-level LRT to the concatenated alignment of a TFBS group, where we estimate two scaling factors, *r*_1_ and *r*_2_, for the foreground and the background branches, respectively. We interpret *r*_1_ and *r*_2_ as the relative substitution rates of the TFBS group in the foreground and background lineages throughout this study. We assume that *r*_1_ = *r*_2_ in the null model (*H*_0_), indicating that the TFBS group has evolved at a constant rate across all lineages. Conversely, we assume that *r*_1_ ≠ *r*_2_ in the alternative model (*H*_*a*_), indicating that the TFBS group has evolved at different substitution rates between the foreground and background lineages. Because the TFBS group may consist of hundreds of TFBSs, we assume that the likelihood ratio statistic of the group-level LRT asymptotically follows a chi-square distribution with one degree of freedom. If the null model is rejected in the group-level LRT and *r*_1_ *> r*_2_, we consider that the TFBS group as a whole may be subject to accelerated evolution.

**Figure 1:**
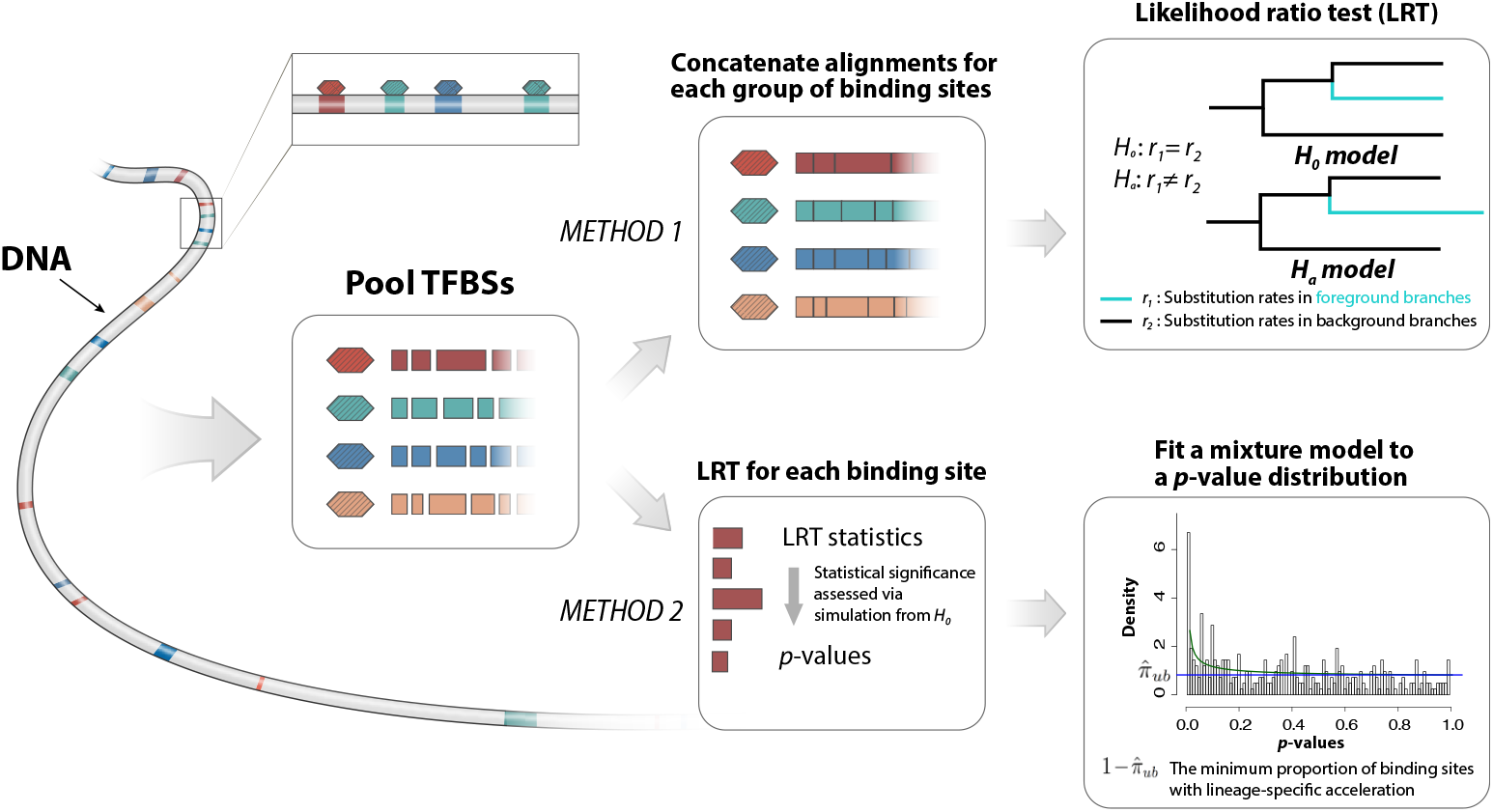
Pooling-based phylogenetic methods for inferring accelerated evolution in TFBSs. In the first method, we fit the group-level LRT to the concatenated alignment of TFBSs bound by the same transcription factor, which allows us to examine whether the group of TFBSs as a whole evolved at an elevated substitution rate in the foreground lineage compared to the background lineage. In the second method, we fit the element-level LRT to the alignment of each individual TFBS, which provides an element-level *p*-value. Then, we fit a beta-uniform mixture model to the distribution of *p*-values in each TFBS group to estimate the proportion of accelerated TFBSs. Colored rectangles and hexagons represent TFBSs and transcription factors, respectively. *r*_1_ and *r*_2_ represent relative substitution rates in the foreground and background lineages, respectively.

In the second method, we use a *phylogenetics-based mixture model* to estimate the proportion of accelerated TFBSs in a TFBS group (Fig. 1). To this end, we first perform an element-level LRT to infer evidence for accelerated evolution in individual TFBSs given that *H*_0_ is rejected in the group-level LRT. The element-level LRT is similar to the group-level LRT but is applied to the alignments of individual TFBSs rather than to the concatenated alignment of the TFBS group. Given the likelihood ratio statistics from the element-level LRT, we then calculate empirical *p*-values for individual TFBSs using parametric bootstrapping. Unlike the chi-square distribution in the group level LRT, the parametric bootstrapping procedure provides accurate *p*-values even when there is a small amount of alignment data per test^9^. Finally, we estimate the proportion of accelerated TFBSs in the TFBS group by fitting a beta-uniform mixture model to the distribution of *p*-values^43^. The beta-uniform mixture model allows us to estimate an upper bound of the proportion of TFBSs generated from *H*_0_ 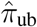. We consider 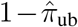 as a conservative estimate (lower bound) of the proportion of accelerated TFBSs.

### Numerous TFBS groups show evidence for accelerated evolution

Using the group-level LRT, we examined accelerated evolution in 4,380,444 TFBSs of 161 transcription factors identified by ChIP-seq experiments in the ENCODE Project^44^. We tested whether each group of TFBSs bound by the same transcription factor had an elevated substitution rate in the human lineage. We used Multiz genome alignments of ten primate species^45^ and defined the human lineage after the divergence of chimpanzees and humans as the foreground lineage. Unlikely previous studies of HARs^7–15^, we did not include non-primate vertebrates to mitigate the impact of the evolutionary turnover of TFBSs on our analysis^30–33, 38, 39^. After Bonferroni correction, we observed that 15 TFBS groups had significantly different substitution rates between the foreground and background lineages (Supplementary Data 1), which all showed elevated substitution rates in humans compared to other primates (*r*_1_ *> r*_2_).

TFBS groups with elevated substitution rates in humans could be either directly under accelerated evolution or merely overlap with other accelerated TFBS groups. To identify TFBS groups directly under accelerated evolution, we sought to partition the binding sites of the 15 TFBS groups with elevated substitution rates into non-overlapping, biologically interpretable TFBS groups. Because BDP1, BRF1, and POLR3G are components of the Pol III transcription machinery^46^, we defined a new TFBS group, Pol III binding, consisting of genomic regions bound by at least two of the three transcription factors. Similarly, since POU5F1 and NANOG can interact with each other to form a protein complex^47, 48^, we defined another TFBS group, POU5F1-NANOG binding, consisting of genomic regions bound by both of the two transcription factors.

Then, we removed all binding sites overlapping more than one TFBS group, resulting in 17 non-overlapping TFBS groups. We applied the group-level LRT again to these non-overlapping TFBS groups. After Bonferroni correction, seven non-overlapping TFBS groups showed significantly elevated substitution rates in the human lineage (Fig. 2; Supplementary Table 1). These non-overlapping TFBS groups included Pol III binding, POU5F1-NANOG binding, BDP1, FOXP2, POU5F1, NANOG, and NRF1. Compared to previously identified HARs, the seven non-overlapping TFBS groups showed weaker acceleration as evidenced by their smaller increases in substitution rates in the human lineage (Fig. 2). We focused on the seven non-overlapping TFBS groups with evidence for accelerated evolution in downstream analysis.

**Figure 2:**
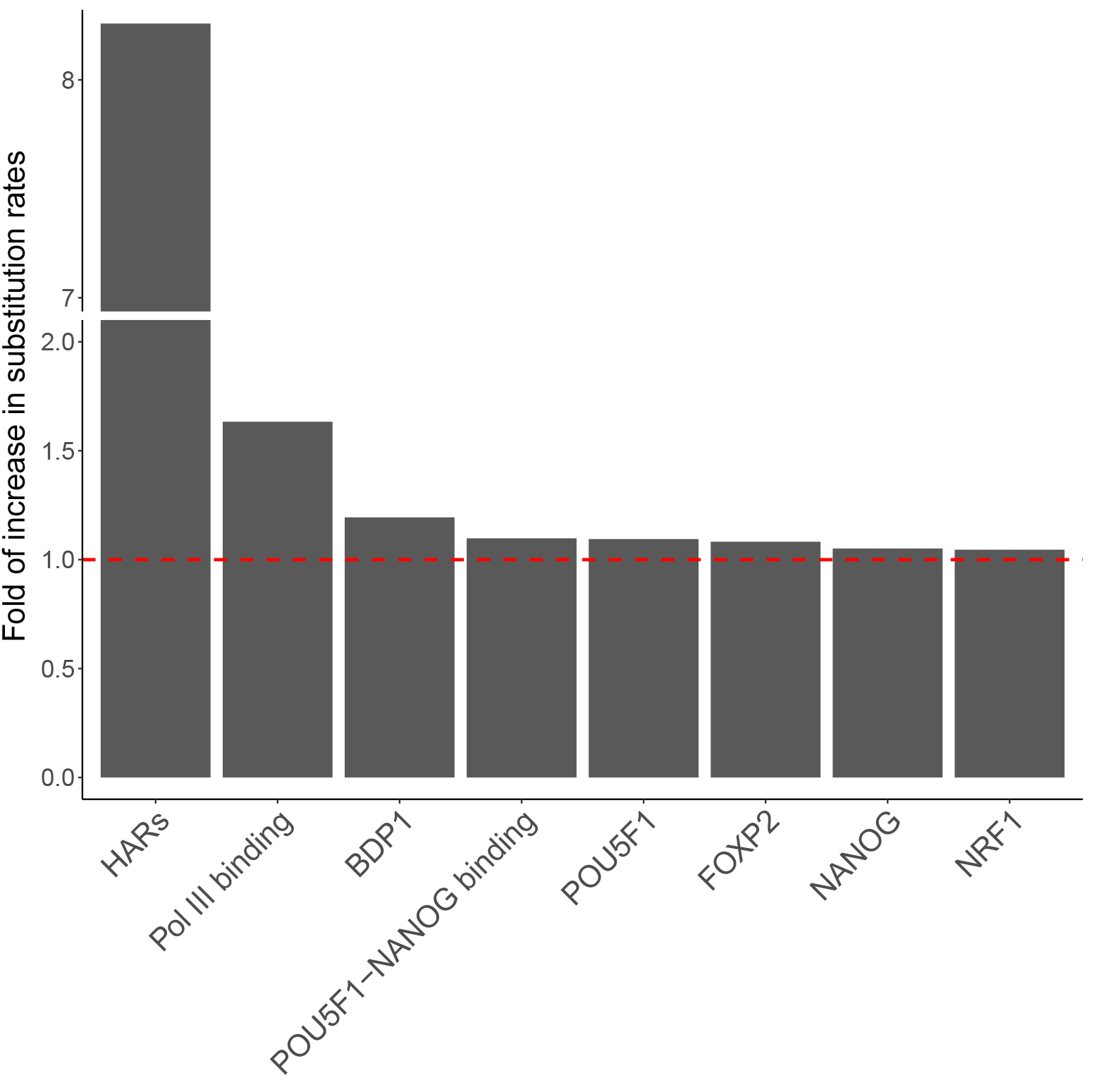
Non-overlapping TFBS groups under accelerated evolution in the human genome. The fold of increase in substitution rate is defined as *r*_1_*/r*_2_, where *r*_1_ and *r*_2_ are the relative substitution rates of a TFBS group in the human lineage and in other primates, respectively.

### Accelerated evolution in TFBSs may not be human specific

A recent study showed that many HARs may also undergo accelerated evolution in other apes^15^. To characterize when acceleration occurred during the evolution of TFBSs, we employed a model comparison approach to search for lineages with elevated substitution rates. Specifically, we evaluated the goodness-of-fit of seven phylogenetic models with different foreground lineages, denoted as M1 to M7 (Fig. 3a). All these models were based on *H*_*a*_ in the group-level LRT (Fig. 1), and the foreground lineages associated with these models corresponded to all the monophyletic clades that included the human lineage (Fig. 3a). These models effectively assumed that the change of substitution rate occurred at most once during the evolution of a TFBS group, which was designed to explore the most parsimonious explanations of accelerated evolution and to limit the number of tested foreground lineages. We used the Bayesian information criterion (BIC) as a measure of goodness-of-fit of these models.

**Figure 3:**
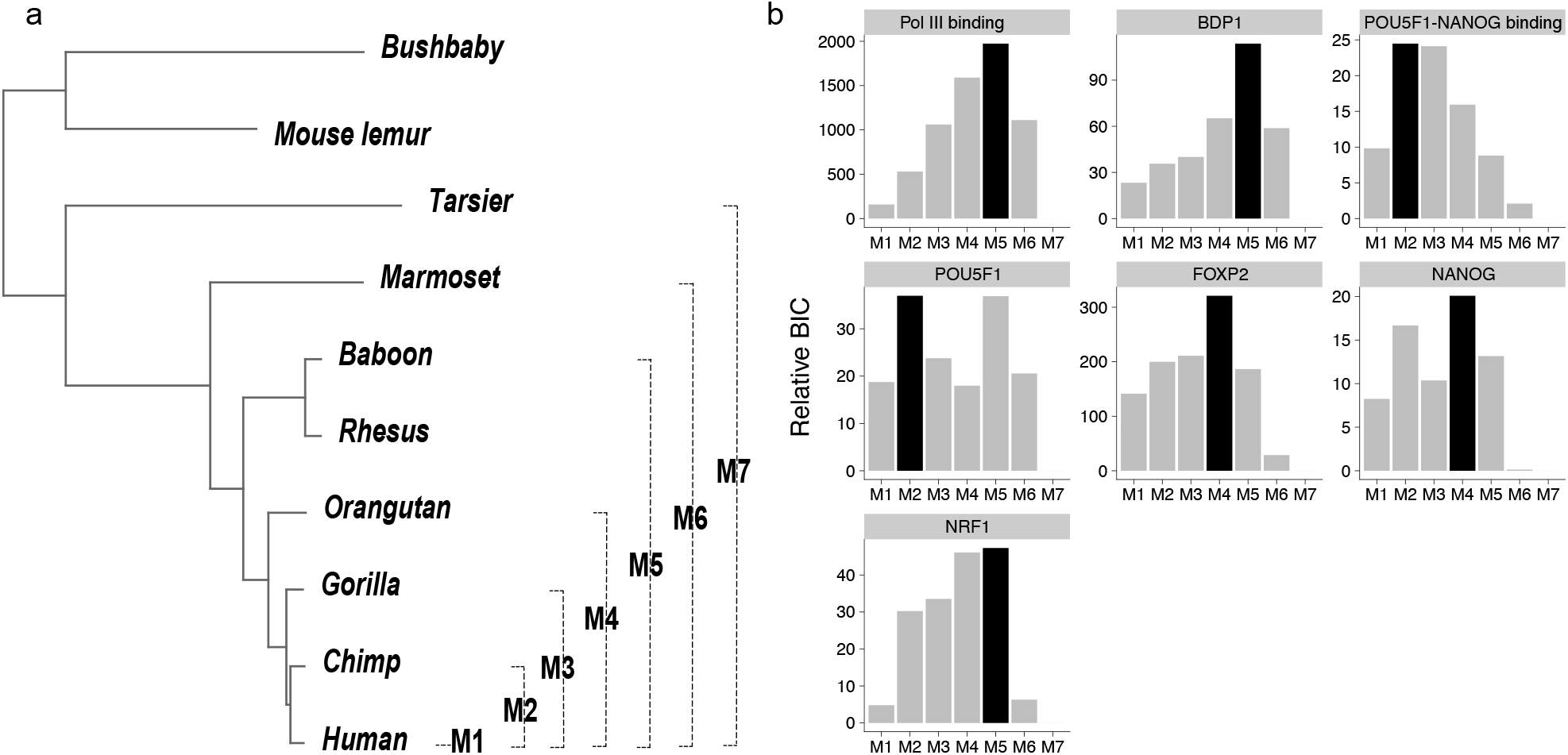
Lineages associated with accelerated evolution in TFBS groups. (a) The seven foreground lineages examined in the model comparison analysis. (b) Model fit with different foreground lineages. The black bars indicate the best-fit foreground lineages.

Although the seven accelerated TFBS groups were originally detected using humans as the foreground lineage, our model comparison analysis showed that accelerated evolution may not be human specific (Fig. 3b, Supplementary Table 2). Specifically, for binding sites of Pol III, BDP1, and NRF1, a model with both apes and Old World monkeys as the foreground lineage (M5) showed the best goodness-of-fit. Similarly, for binding sites of FOXP2 and NANOG, a model with apes as the foreground (M4) showed the best goodness-of-fit. Moreover, for POU5F1-NANOG and POU5F1 binding sites, a model with Hominini as the foreground (M2) showed the best goodness-of-fit. Altogether, the acceleration of TFBS evolution might be driven by changes of selection pressure in Hominini, apes, and Old World monkey and, thus, might contribute to phenotypic differences between these species and other primates.

### More than 6,000 TFBSs may be under accelerated evolution

In this section, we sought to infer the total number of TFBSs under accelerated evolution. While the group-level LRT can examine whether a TFBS group as a whole was under accelerated evolution, it could not estimate the number of accelerated TFBSs in the TFBS group. Also, because the signal of acceleration might be weak in TFBSs (Fig. 2), previous phylogenetic models could not be used to estimate this number either^7–15^. To address this problem, we utilized the phylogenetics-based mixture method to estimate the proportion of accelerated TFBSs from the distribution of *p*-values associated with individual TFBSs in the same group (Fig. 1).

We observed that 78% Pol III binding sites were under accelerated evolution (Table 1), which translates to approximately 222 accelerated Pol III binding sites. Also, 20% and 25% binding sites of BDP1 and NRF1 were under accelerated evolution in Old World monkeys and apes, which translate to approximately 90 and 466 accelerated TFBSs, respectively. Approximately 25% TFBSs of FOXP2 and NANOG were under accelerated evolution in apes, suggesting about 5,000 binding sites in these TFBS groups were accelerated elements. Furthermore, approximately 8% TFBSs of POU5F1 and POU5F1-NANOG were under accelerated evolution in Hominini, indicating that about 300 binding sites of the two groups were accelerated in the clade consisting of humans and chimpanzees. In total, more than 6,000 TFBSs were under accelerated evolution in Hominini, apes, and Old World monkeys (Table 1), which is more more than the 3098 known HARs (See Materials and methods).

**Table 1:**
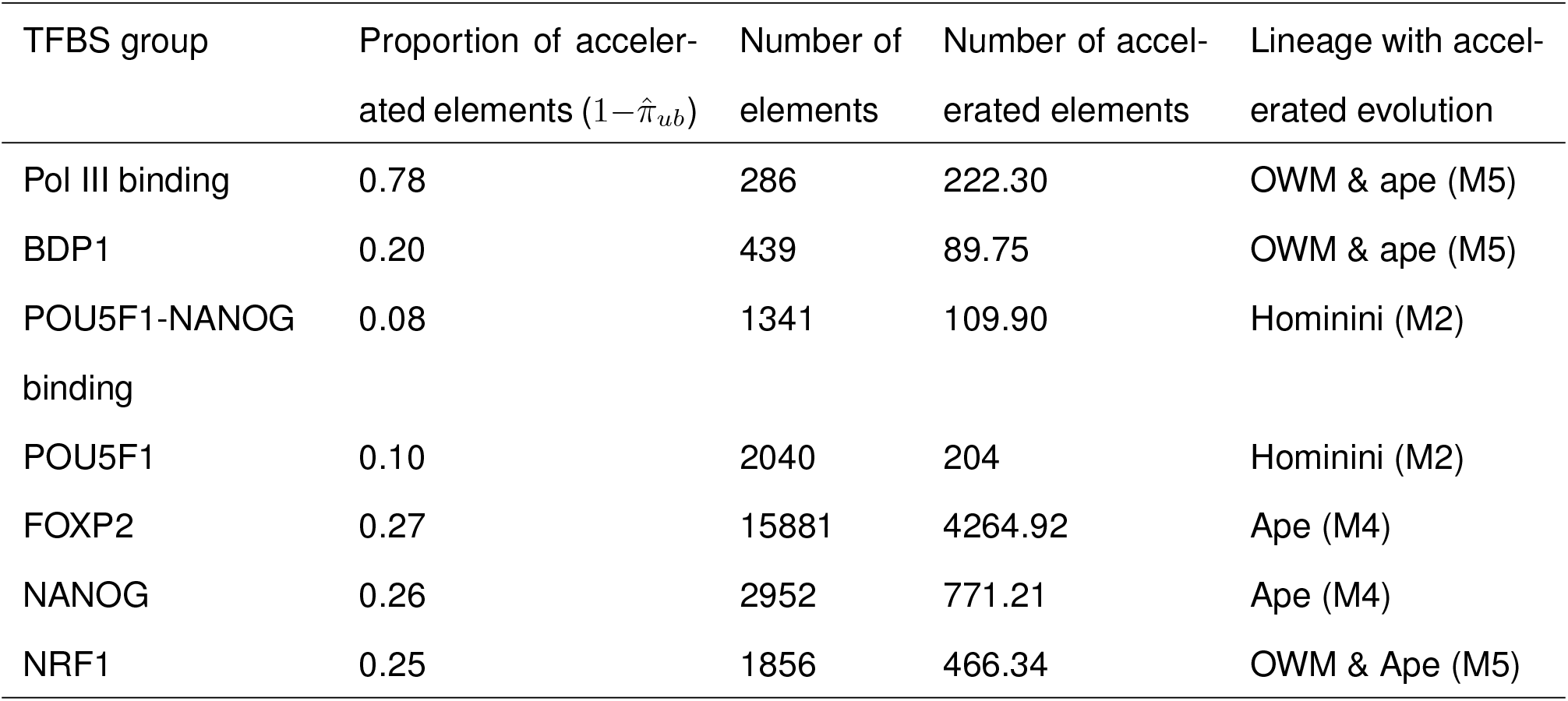
Numbers of accelerated TFBSs estimated by the phylogenetic mixture model.

### Positive selection may drive accelerated evolution in Pol III binding sites

The acceleration of TFBS evolution could be due to either positive selection or relaxed purifying selection in the foreground lineage. To examine whether positive selection is a driver of accelerated evolution in TFBSs, we employed the INSIGHT model^49–51^ to infer the strength of positive selection on the seven accelerated TFBS groups in the human lineage. Similar to the McDonald-Kreitman test^52, 53^, INSIGHT incorporates divergence and polymorphism data to infer positive selection on a set of predefined genomic elements. We fit the INSIGHT model to the binding sites of each TFBS group, which provided an estimate of *D*_*p*_, that is, the expected number of adaptive substitutions per kilobase in the human lineage.

We observed that Pol III binding sites were subject to strong positive selection in the human lineage, because *D*_*p*_ of Pol III binding sites was significantly higher than 0 and was comparable to that of previously identified HARs (Fig. 4; Supplementary Table 3). In other TFBS groups, *D*_*p*_ was not significantly different from 0, indicating that positive selection might not be the driving force of accelerated evolution in these TFBS groups.

**Figure 4:**
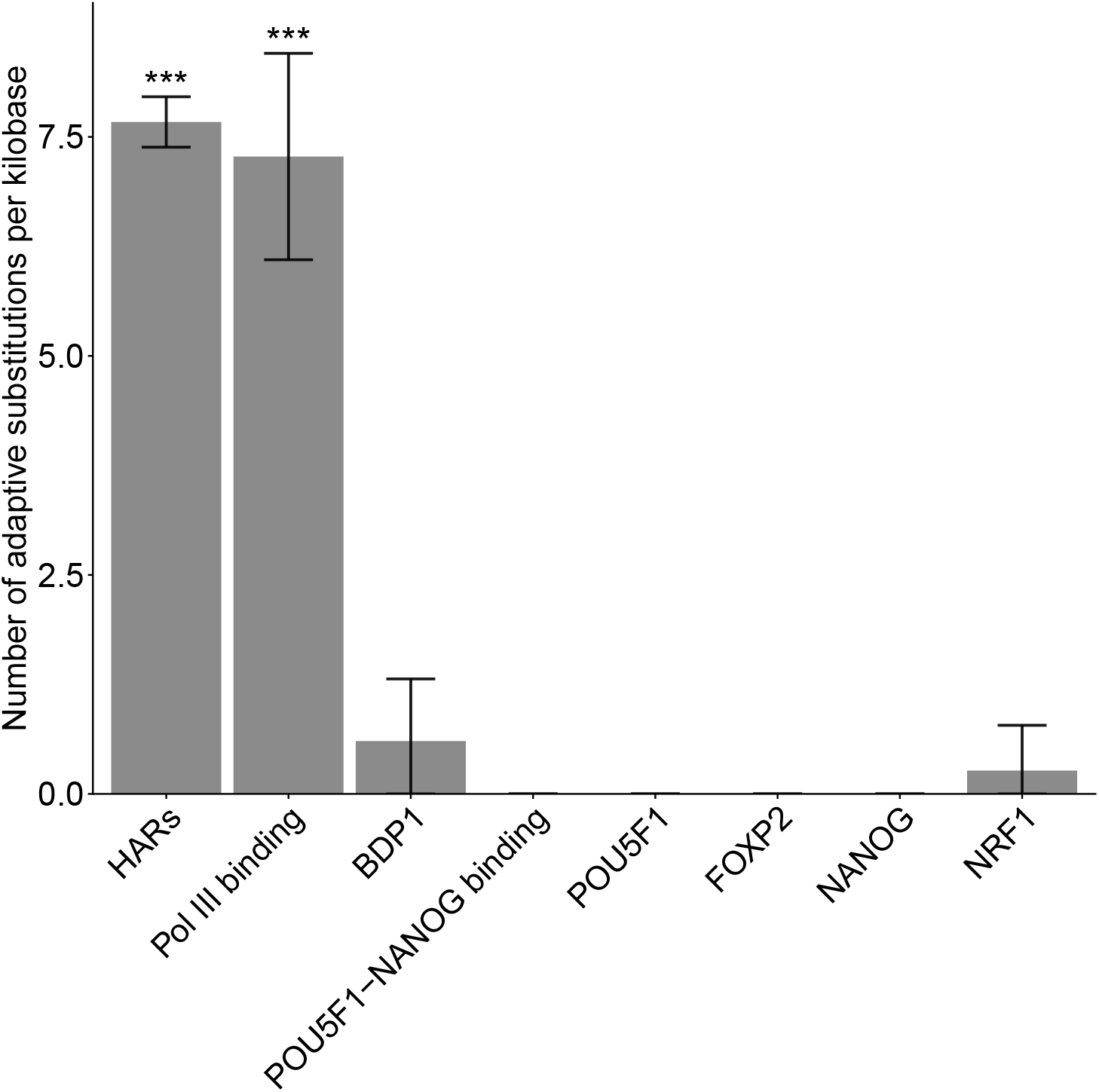
Positive selection on accelerated TFBS groups in the human lineage. The numbers of adaptive substitutions per kilobase are estimated by INSIGHT^49–51^. Error bars represent two-fold standard errors.

### The accelerated TFBSs are enriched around neurodevelopmental and pluripotency genes

To identify the major functions represented by the accelerated TFBS groups, we utilized Genomic Regions Enrichment of Annotations Tool (GREAT) to first find the potential target genes by predicting both proximal and distal binding events, and then analyze the functional significance of those TFBS groups by applying GO enrichment test and pathway enrichment analysis to the potential target genes^54–56^. Using default settings in GREAT, we found 9896 potential target genes for FOXP2 binding sites, 3745 genes for NANOG binding sites, 2931 genes for NRF1 binding sites, 1976 genes for POU5F1-NANOG binding sites and 478 potential target genes for POU5F1 binding sites. After GO enrichment test and removing the redundant GO terms with high semantic similarity (0.7) and performing bonferroni correction, we found FOXP2 binding sites were enriched in 178 biological process(BP) GO terms including developmental processes (such as forebrain development) and neural system (Fig. 5a). NANOG binding sites were enriched in 83 BP GO terms mainly about development (Fig. 5b). NRF1 binding sites were enriched in 7 GO terms including ear development and chromatin modification(Fig. 5c). POU5F1-NANOG complex were enriched in 52 BP GO terms related to organization and regulation of synapse (Fig. 5d). To investigate the functions of TFBSs at pathway level, the ReactomePA package ^56^ was used to analyze the potential target genes of each TFBS group. The potential target genes of NANOG binding sites and POU5F1-NANOG binding sites were enriched in the pathways involved with transcriptional regulation of pluripotent stem cells(Fig. 6a,b). FOXP2 binding sites were shown to be associated with multiple signaling pathways(Fig. 6c). In the genes associated with other TFBS groups, no pathways or biological terms were found to be significant after correction.

**Figure 5:**
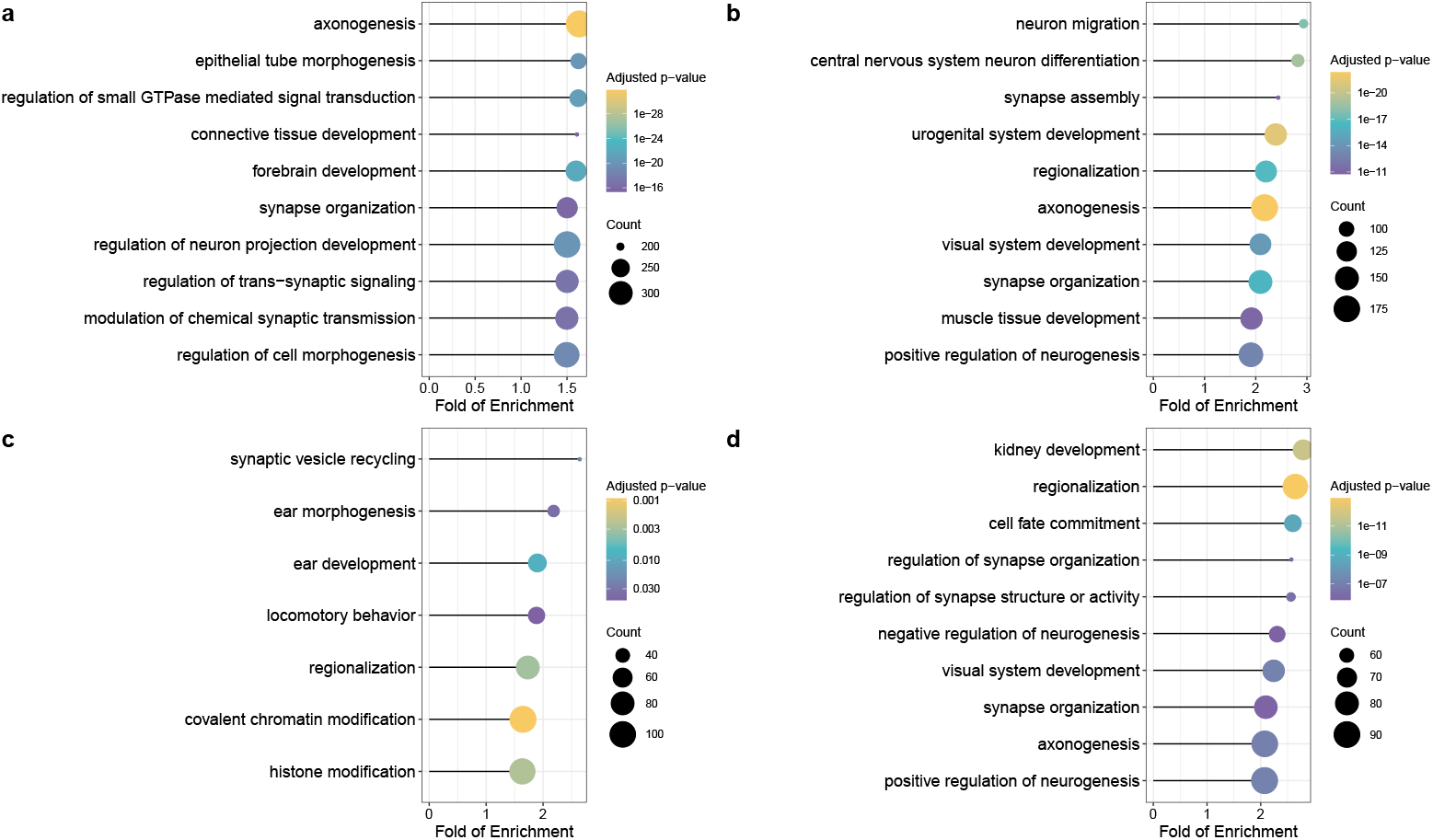
Gene ontology analysis of the accelerated TFBS associated genes. The dot plots show the top 10 significant GO terms for biological process of (a) FOXP2 binding sites associated genes (b) NANOG binding sites associated genes (c) NRF1 binding sites associated genes (d) POU5F1-NANOG binding associated genes. The x-axis represents the fold of enrichment. The size of circle represents the number of TFBS-associated genes affiliated with the specific GO terms. The color of circle represents the Bonferroni-corrected *p*-values.

**Figure 6:**
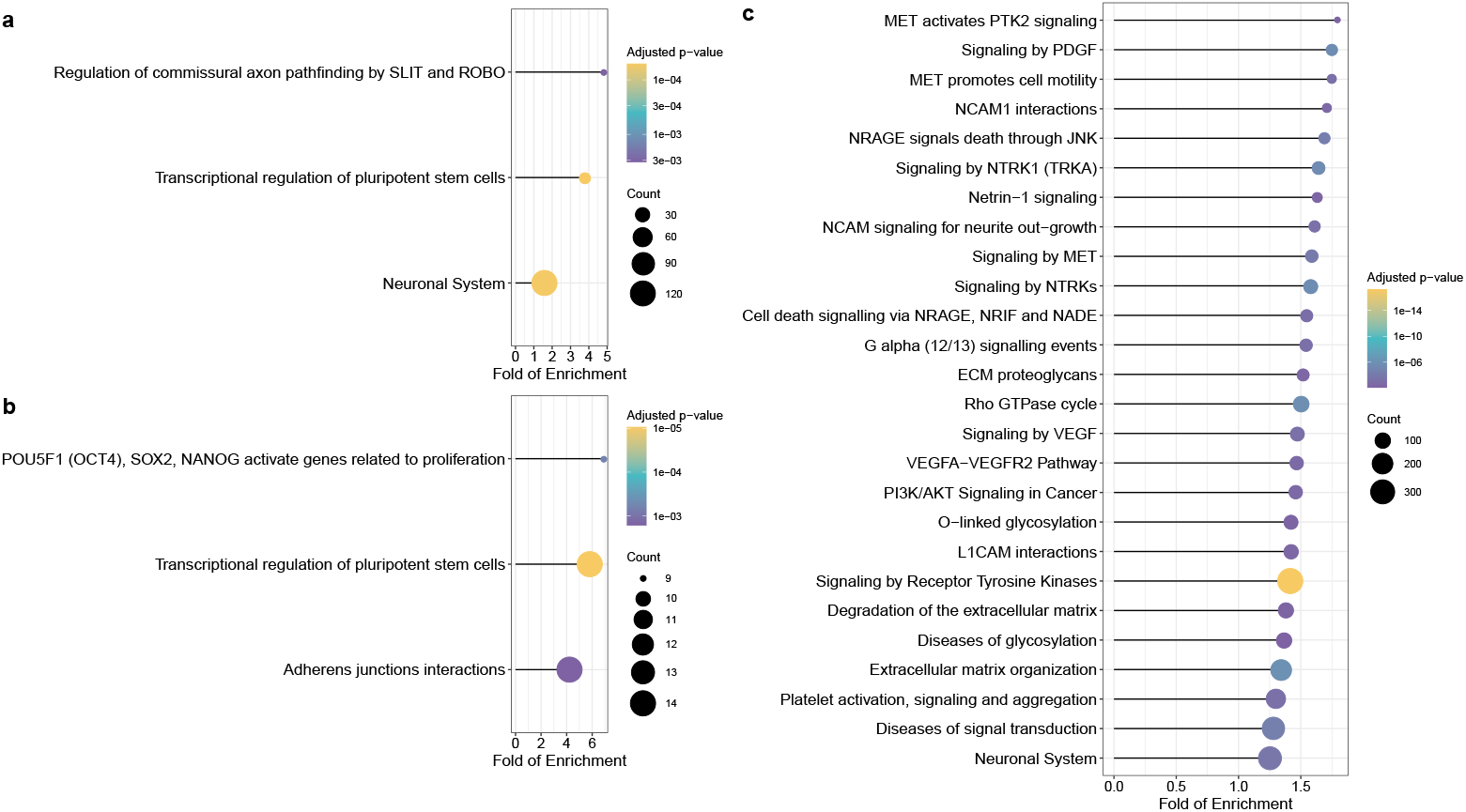
Reactome pathway analysis of the accelerated TFBS associated genes. The dot plots showing the significant pathways after Bonferroni correction of (a) NANOG binding sites’ associated genes (b) POU5F1-NANOG binding sites’ associated genes (c) FOXP2 binding sites associated genes. The x-axis represents the fold of enrichment. The size of circle represents the number of TFBS-associated genes affiliated with the specific GO terms. The color of circle represents the Bonferroni-corrected *p*-values.

## Discussion

In the current study, we present two pooling-based methods to infer genomic elements under accelerated evolution. Unlike previous methods that focus on analyzing individual elements^7–15^, our new methods group hundreds of genomic elements with similar biological functions to increase the sample size per test and reduce the multiple testing burden. Thus, our methods may have higher sensitivity to detect weak signals of accelerated evolution. To the best of our knowledge, our methods are the first statistical framework dedicated to infer weakly accelerated evolution.

Using the group-level LRT, we identify seven groups of non-overlapping TFBSs with significant evidence for accelerated evolution (Fig. 2). The model comparison analysis suggests that these TFBS groups may be under accelerated evolution not only in humans but also in other primate species (Fig. 3). In agreement with our finding, a recent study of HARs has shown that many HARs may also be subject to accelerated evolution in other ape species^15^. Therefore, accelerated evolution of regulatory elements may be a shared characteristic of primates rather than specific to the human lineage.

Among the seven groups of accelerated TFBSs, we show that Pol III binding sites may be subject to positive selection in the human lineage but find no evidence for positive selection in other accelerated TFBS groups (Fig. 4). Therefore, both lineage-specific positive selection and nonadaptive evolutionary forces, such as relaxed purifying selection and GC-biased gene conversion^21, 57^, may drive the accelerated evolution of TFBSs. In contrast, more than half of HARs may be subject to positive selection in the human lineage^21^, suggesting that positive selection may be the main driving force of accelerated evolution in HARs. Because previous studies of HARs have focused on identifying individual genomic elements with extremely high substitution rates in the human lineage, the higher frequency of detecting positive selection in HARs could partially reflect the lower power of previous methods in discovering weakly accelerated elements driven by evolutionary forces other than positive selection.

Although accelerated evolution is much weaker in Pol III binding sites than in HARs (Fig. 2), Pol III binding sites are subject to strong positive selection in the human lineage, on par with HARs (Fig. 4). Because HARs are highly conserved across species, they may have very low substitution rates in non-human primates, which in turn enhances the signals of accelerated evolution. In contrast, Pol III binding sites may not be highly conserved across species, resulting in a weaker signal of accelerated evolution despite strong positive selection in the foreground lineage. Taken together, weak signals of accelerated evolution may not always imply weak positive selection in the foreground lineage.

Notably, the seven groups of accelerated TFBSs may play key roles in developmental processes. First, recent studies suggest that disruptive mutations in subunits of Pol III, such as *POLR3A, POLR3B* and *BRF1*, may be associated with neurodevelopmental disorders^58–60^. Therefore, accelerated evolution in Pol III binding sites might be associated with the adaptive evolution of the central nervous system in apes and Old World monkeys (Fig. 3). Second, POU5F1 and NANOG are transcription factors necessary to the pluripotency and self-renewal of embryonic stem cells^61–63^. The colocalization of POU5F1 and NANOG in regulatory elements, referred to as POU5F1-NANOG binding in the current study, might trigger zygotic gene activation in vertebrates^47, 64–67^. Third, FOXP2 is a highly conserved vertebrate protein with high expression in the central nervous systems during embryogenesis, and detrimental mutations in the *FOXP2* gene may cause impaired speech development in humans^68–70^. Also, previous studies have shown that the protein sequence and expression of *FOXP2* could be subject to accelerated evolution in humans^71–73^, echolocating bats^74^, and vocal learning birds^75^. Finally, NRF1 has been found to regulate the expression of *GABRB1*, a gene associated with neurological and neuropsychiatric disorders^76, 77^. Together with the fact that a large proportion of HARs are neural enhancers and subject to accelerated evolution in humans and other primates^19–21^, we conclude that regulatory sequences of neurodevelopmental genes may be the main target of accelerated evolution in primates.

Due to the scarcity of ChIP-seq data in non-human primates, we have used human-based TFBS annotations to infer accelerated evolution. It may limit our ability to detect accelerated evolution present in non-human primates but not in humans. Thus, our estimate of the number of accelerated TFBSs is likely to be conservative (Table 1). In future studies, it is of great interest to investigate accelerated evolution in TFBSs identified in non-human primates, highlighting the urgent need to perform high-throughput functional genomic experiments in our close relatives.

Compared to conserved genomic elements explored in previous studies of HARs, TFBSs may have a higher evolutionary turnover rate^30–33, 38, 39^. To alleviate the impact of evolutionary turnover on our analysis, we have only included primate genomes in the current study. Nevertheless, a small proportion of TFBSs identified in the human genome may still be subject to evolutionary turnover in other primates^39^. We expect that the evolutionary turnover of TFBSs in non-human primates may not lead to false positive results in our analysis. Indeed, conditional on the presence of a TFBS in the human genome, the evolutionary turnover of the TFBS in non-human primates is more likely to increase the substitution rate in the background lineage and hence makes our analysis conservative. Once ChIP-seq data become available in multiple non-human primates in the future, the conservativeness of our analysis may be alleviated by including only species where the TFBS of interest is detected.

Our pooling-based methods have a potential to be extended in future studies. For instance, if multiple TFBSs overlap with each other, our current methods cannot distinguish between TFBSs directly under accelerated evolution from those overlapping other accelerated TFBSs. To address this problem, we have used a heuristic method to remove overlapping TF-BSs in the current study, which may reduce the number of TFBSs in our analysis. In the future, it is of great interest to develop a rigorous method for inferring accelerated evolution in overlapping TFBSs. Motivated by the recent success of evolution-based regression models^34, 78–80^, we propose that unifying our pooling-based methods and generalized linear models may be a promising direction to disentangle causal from correlational relationships in the analysis of accelerated evolution.

## Methods

### Genome alignment and TFBS annotation

We obtained the Multiz alignment of 46 vertebrate genomes from the UCSC Genome Browser^45^. Then, we extracted a subset of alignments for ten primate species from the 46-way Multiz alignment. The ten primate species and their genome assemblies included *Homo Sapiens* (hg19), *Pan troglodytes* (panTro2), *Gorilla gorilla* (gorGor1), *Pongo abelii* (ponAbe2), *Macaca mulatta* (rheMac2), *Papio hamadryas* (papHam1), *Callithrix jacchus* (calJac1), *Tarsius syrichta* (tarSyr1), *Microcebus murinus* (micMur1), and *Otolemur garnettii* (otoGar1). Also, we downloaded 4,380,444 TFBSs for 161 transcription factors from the UCSC Genome Browser. These TFBSs were identified by ChIP-seq experiments in the ENCODE Project^44^. We extracted alignments of TFBSs across ten primate species using PHAST^81^. We removed the TFBSs overlapping with UTRs, CDSs, and previously identified HARs. To filter out low-quality alignments, we obtained informative alignment sites where unambiguous bases were found in at least five out of ten primate species in the Multiz alignment. We retained TFBSs with at least 50 informative alignment sites for downstream analysis.

### Previously defined HARs collection

We obtained a comprehensive list of previously defined HARs from https://docpollard.org/research/. We first combined the following genetic elements: Merged list of 2649 HARs(a set of HARs in noncoding regions built by Capra et al.^17^), 284 human accelerated elements in mammal conserved regions with adjusted *p*-value *<*0.05 (mapped to hg19 using the LiftOver tools on the UCSC genome browser), and 760 human accelerated elements in primate conserved regions with adjusted *p*-value *<*0.05 (mapped to hg19 using the LiftOver tools on the UCSC genome browser). Then we sorted and merged the bed file using bedtools/2.27.1.

### Group-level LRT for inferring accelerated evolution

We built a reference phylogenetic model using the alignment of ten primate genomes, assuming that the majority of TFBSs may not be subject to accelerated evolution. We first concatenated alignments of all TFBSs. We then fit a phylogenetic model to the concatenated alignment using the phangorn library in R^82^. In the phylogenetic model, we used the generalized time-reversible (GTR) substitution model to describe nucleotide sequence evolution and the discrete Gamma distribution with four rate categories to model substitution rate variation among nucleotide sites^41^. Also, we fixed the tree topology of the reference phylogenetic model to the one used from the UCSC Genome Browser (http://hgdownload.cse.ucsc.edu/goldenPath/hg19/multiz46way/46way.nh). We estimated model parameters, including branch lengths, the shape parameter of the discrete Gamma distribution, and parameters of the GTR substitution model, using the optim.pml function in phangorn.

Given the reference phylogenetic model, we used a customized R program based on phangorn to perform the group-level LRT. First, we concatenated alignments for each TFBS group separately. Then, we fit two group-level phylogenetic models to the concatenated alignment of each TFBS group. In the null model (*H*_0_), we inferred a global scaling factor of branch lengths with maximum likelihood estimation and fixed all other model parameters to the ones in the reference phylogenetic model. We interpreted the estimated scaling factor as the relative substitution rate of TFBS sequences in both the foreground and the background lineage. In the alternative model (*H*_*a*_), we estimated two scaling factors of branch lengths, *r*_1_ and *r*_2_, for the foreground and background lineages, respectively. The two scaling factors were interpreted as the relative substitution rates in the foreground and background lineages in the alternative model. For each TFBS group, we calculated a likelihood ratio statistic defined as the two-fold difference in the log likelihood between *H*_*a*_ and *H*_0_. Given the likelihood ratio statistic, we obtained a *p*-value for each TFBS group using a chi-square test with one degree of freedom. Finally, we calculated adjusted *p*-values using the Bonferroni correction.

From the group-level LRT, we found that TFBSs of 15 transcription factors showed elevated substitution rates in the human lineage. We further partitioned the TFBSs of the 15 transcription factors into 17 non-overlapping TFBS groups. These non-overlapping TFBS groups included genomic regions exclusively bound by one of the 15 transcription factors and two new TFBS groups: Pol III binding and POU5F1-NANOG binding. The Pol III binding group consisted of TFBSs bound by at least two of BDP1, BRF1, and POLR3G. Similarly, the group of POU5F1-NANOG binding consisted of TFBSs bound by both POU5F1 and NANOG. Then, we applied the group-level LRT again to the 17 non-overlapping TFBS groups and calculated adjusted *p*-values using the Bonferroni correction.

### Reduction of redundancy in the 15 TFBS groups

From Group-level LRT, we found 15 groups of TFBSs with accelerated evolution in human. However, there is redundancy among the data possibly because the transcription factors share a considerable proportion of binding sites.

Some of the groups have similar biological functions, for example, BDP1, BRF1 and POLR3G are key factors in the Pol III transcription machinery; POU5F1 and NANOG are necessary regulators in ES cell pluripotency and self-renewal. To identify the evolutionary forces in the colocalization of transcription factors, we defined two new TFBS groups. The Pol III binding sites, were defined as the binding sites occupied by at least two out of the three transcription factors related to Pol III (BDP1, BRF1 and POLR3G). To define the POU5F1-NANOG binding, we obtained the intersecting regions of POU5F1 and NANOG binding sites.

To remove redundancy in overlapping binding sites, we then got the non-overlapping regions bound by merely BDP1, BRF1 or POLR3G. For each of the other 12 TFBS groups with accelerated evolution in human, we obtained the entries that don’t overlap with any of BDP1, BRF1, POLR3G or the rest 11 TFBS groups. Then we ran the group-level LRT again for the 15 non-overlapping TFBS groups and 2 newly-defined TFBS groups.

### Inference of lineages with accelerated evolution

We utilized the alternative model (*H*_*a*_) in the group-level LRT to search for lineages associated with accelerated evolution. To this end, we fit the group-level *H*_*a*_ with seven different foreground lineages to the concatenated alignment of each TFBS group (Fig. 3). The seven foreground lineages corresponded to all monophyletic clades that included humans. For each TFBS group and foreground lineage, we used the BIC as a measure of goodness-of-fit,

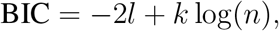

where *l* is the log likelihood of the group-level *H*_*a*_, *k* is the number of model parameters, and *n* is the sample size. Because the group-level *H*_*a*_ included two parameters (*r*_1_ and *r*_2_), we set *k* to 2. Also, we assumed that *n* could be approximated by the total number of bases in the concatenated alignment of each TFBS group. For each TFBS group, we considered the foreground lineage with the highest BIC to be the best-fit lineage.

### Estimation of the number of TFBSs under accelerated evolution

We utilized the R program for the group-level LRT to perform the element-level LRT. To this end, we applied the R program to the alignment of each individual TFBS separately, after filtering out TFBSs with less than 50 informative alignment sites. Then, we performed parametric bootstrapping at the group level to calculate a *p*-value for each TFBS. Specifically, we first fit the *H*_0_ in the group-level LRT to the concatenated alignment of each TFBS group, which provided a global scaling factor to calibrate the branch lengths of the reference phylogenetic model. Second, we randomly sampled 10,000 TFBSs with replacement from each TFBS group and used the calibrated phylogenetic model to generate 10,000 simulated alignments of matched length. Third, we fit the element-level LRT to the simulated TFBS alignments from the same group, which provided an empirical null distribution of the likelihood ratio statistic for each TFBS group. Fourth, we compared the observed likelihood ratio statistic to the empirical null distribution to calculate a *p*-value for each TFBS. Finally, we fit a beta-uniform mixture model to *p*-values from each TFBS group^43^. We considered a statistic, 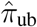, from the beta-uniform mixture model as the proportion or binding site without acceleration and, accordingly, 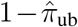 as the proportion of accelerated TFBSs.

### Detection of selection pressure

To investigate if accelerated evolution in TFBSs were driven by positive selection, we used the INSIGHT model to infer positive selection on the seven accelerated, non-overlapping TFBS groups in the human lineage^49–51^. We obtained INSIGHT2, a highly efficient implementation of the INSIGHT model, from https://github.com/CshlSiepelLab/FitCons2. Then, we applied INSIGHT2 to each TFBS group. INSIGHT2 provided *D*_*p*_ and SE[*D*_*p*_], that is, the expected number of adaptive substitutions per kilobase and its standard error. We performed the Wald test to examine if *D*_*p*_ was significantly different from 0 for each TFBS group. Under the null hypothesis of *D*_*p*_ = 0, we assumed that the z-statistic, 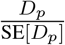, asymptotically followed a 50:50 mixture of a point probability mass at 0 and a half standard normal distribution^83^.

### Functional enrichment analysis of accelerated TFBS associated genes

We identified the potential target genes of accelerated TFBSs using GREAT with default settings. Then we perfomed GO enrichment analysis using clusterProfiler and ran pathway enrichment analysis using ReactomePA. With the clusterProfiler package, the significance of enrichment test for GO terms under biology process subontology was estimated by hypergeometric distribution and then adjusted by Bonferroni correction. The redundant GO terms were trimmed by applying the *simplify* to remove terms among which semantic similarities were higher than 0.7. Only top ten genes with highest folds of enrichments were shown in the Fig. 5, while the complete list of significant GO biological process terms with corrected *p*-value *<*0.05 are available in the Supplementary Data 2, With the ReactomePA package, we first estimated the significance by hypergeometric distribution and then applied Bonferroni correction to the results. The complete list of significant pathways with corrected *p*-value *<*0.05 are available in the Supplementary Data 2.

## Data Availability

GroupAcc and companion data are available at https://github.com/yifei-lab/GroupAcc/.

## Acknowledgement

The authors thank Bohao Fang, Adam Siepel, Ilan Gronau, Zhihan Liu and Ritika Ramani for useful discussions. Research reported in this publication was supported by the National Institute of General Medical Sciences of the National Institutes of Health under Award Number R35GM142560 and by the Pennsylvania State University. The content is solely the responsibility of the authors and does not necessarily represent the official views of the National Institutes of Health.

## Notes

### Competing Interest Statement

The authors have declared no competing interest.

## References

1. Haygood, R., Fedrigo, O., Hanson, B., Yokoyama, K.-D. & Wray, G. A. Promoter regions of many neural- and nutrition-related genes have experienced positive selection during human evolution. Nature Genetics 39, 1140 (2007).

2. Kosiol, C. et al. Patterns of positive selection in six mammalian genomes. PLOS Genetics 4, e1000144 (2008).

3. Sackton, T. B. et al. Convergent regulatory evolution and loss of flight in paleognathous birds. Science 364, 74–78 (2019).

4. Zhao, S. et al. Identifying lineage-specific targets of natural selection by a bayesian analysis of genomic polymorphisms and divergence from multiple species. Molecular Biology and Evolution 36, 1302–1315 (2019).

5. Clark, A. G. et al. Inferring nonneutral evolution from human-chimp-mouse orthologous gene trios. Science 302, 1960–1963 (2003).

6. Dorus, S. et al. Accelerated evolution of nervous system genes in the origin of Homo sapiens. Cell 119, 1027–1040 (2004).

7. Prabhakar, S., Noonan, J. P., Pääbo, S. & Rubin, E. M. Accelerated evolution of conserved noncoding sequences in humans. Science 314, 786–786 (2006).

8. Pollard, K. S. et al. An RNA gene expressed during cortical development evolved rapidly in humans. Nature 443, 167–172 (2006).

9. Pollard, K. S. et al. Forces shaping the fastest evolving regions in the human genome. PLOS Genetics 2, e168 (2006).

10. Kim, S. Y. & Pritchard, J. K. Adaptive evolution of conserved noncoding elements in mammals. PLOS Genetics 3, e147 (2007).

11. Bird, C. P. et al. Fast-evolving noncoding sequences in the human genome. Genome Biology 8, R118 (2007).

12. Bush, E. C. & Lahn, B. T. A genome-wide screen for noncoding elements important in primate evolution. BMC Evolutionary Biology 8, 17 (2008).

13. Lindblad-Toh, K. et al. A high-resolution map of human evolutionary constraint using 29 mammals. Nature 478, 476–482 (2011).

14. Gittelman, R. M. et al. Comprehensive identification and analysis of human accelerated regulatory DNA. Genome Research 25, 1245–1255 (2015).

15. Kostka, D., Holloway, A. K. & Pollard, K. S. Developmental loci harbor clusters of accelerated regions that evolved independently in ape lineages. Molecular Biology and Evolution 35, 2034–2045 (2018).

16. Prabhakar, S. et al. Human-specific gain of function in a developmental enhancer. Science 321, 1346–1350 (2008).

17. Capra, J. A., Erwin, G. D., Gabriel, M., Rubenstein, J. L. R. & Pollard, S., Katherine. Many human accelerated regions are developmental enhancers. Philosophical Transactions of the Royal Society B: Biological Sciences 368, 20130025 (2013).

18. Kamm, G. B., Pisciottano, F., Kliger, R. & Franchini, L. F. The developmental brain gene npas3 contains the largest number of accelerated regulatory sequences in the human genome. Molecular Biology and Evolution 30, 1088–1102 (2013).

19. Ryu, H. et al. Massively parallel dissection of human accelerated regions in human and chimpanzee neural progenitors. bioRxiv (2018).

20. Uebbing, S. et al. Massively parallel discovery of human-specific substitutions that alter enhancer activity. Proceedings of the National Academy of Sciences 118 (2021).

21. Kostka, D., Hubisz, M. J., Siepel, A. & Pollard, K. S. The role of GC-biased gene conversion in shaping the fastest evolving regions of the human genome. Molecular Biology and Evolution 29, 1047–1057 (2012).

22. Xu, K., Schadt, E. E., Pollard, K. S., Roussos, P. & Dudley, J. T. Genomic and network patterns of schizophrenia genetic variation in human evolutionary accelerated regions. Molecular Biology and Evolution 32, 1148–1160 (2015).

23. Doan, R. N. et al. Mutations in human accelerated regions disrupt cognition and social behavior. Cell 167, 341–354 (2016).

24. Levchenko, A., Kanapin, A., Samsonova, A. & Gainetdinov, R. R. Human accelerated regions and other human-specific sequence variations in the context of evolution and their relevance for brain development. Genome Biology and Evolution 10, 166–188 (2018).

25. Wei, Y. et al. Genetic mapping and evolutionary analysis of human-expanded cognitive networks. Nature Communications 10, 4839 (2019).

26. Castelijns, B. et al. Hominin-specific regulatory elements selectively emerged in oligodendrocytes and are disrupted in autism patients. Nature Communications 11, 301 (2020).

27. Booker, B. M. et al. Bat accelerated regions identify a bat forelimb specific enhancer in the hoxd locus. PLOS Genetics 12, 1–21 (2016).

28. Eckalbar, W. L. et al. Transcriptomic and epigenomic characterization of the developing bat wing. Nature Genetics 48, 528–536 (2016).

29. Tollis, M. et al. Elephant genomes reveal accelerated evolution in mechanisms underlying disease defenses. Molecular Biology and Evolution 38, 3606–3620 (2021).

30. Dermitzakis, E. T. & Clark, A. G. Evolution of transcription factor binding sites in mammalian gene regulatory regions: Conservation and turnover. Molecular Biology and Evolution 19, 1114–1121 (2002).

31. Moses, A. M. et al. Large-scale turnover of functional transcription factor binding sites in drosophila. PLOS Computational Biology 2, e130 (2006).

32. Doniger, S. W. & Fay, J. C. Frequent gain and loss of functional transcription factor binding sites. PLOS Computational Biology 3, e99 (2007).

33. Schmidt, D. et al. Five-vertebrate ChIP-seq reveals the evolutionary dynamics of transcription factor binding. Science 328, 1036–1040 (2010).

34. Dukler, N., Huang, Y.-F. & Siepel, A. Phylogenetic modeling of regulatory element turnover based on epigenomic data. Molular Biology and Evolution 37, 2137–2152 (2020).

35. Wittkopp, P. J. & Kalay, G. Cis-regulatory elements: molecular mechanisms and evolutionary processes underlying divergence. Nature Reviews Genetics 13, 59–69 (2012).

36. Siepel, A. & Arbiza, L. Cis-regulatory elements and human evolution. Current Opinion in Genetics & Development 29, 81 – 89 (2014).

37. Villar, D., Flicek, P. & Odom, D. T. Evolution of transcription factor binding in metazoans: mechanisms and functional implications. Nature Reviews Genetics 15, 221–233 (2014).

38. Rands, C. M., Meader, S., Ponting, C. P. & Lunter, G. 8.2% of the human genome is constrained: variation in rates of turnover across functional element classes in the human lineage. PLOS genetics 10, e1004525 (2014).

39. Yokoyama, K. D., Zhang, Y. & Ma, J. Tracing the evolution of lineage-specific transcription factor binding sites in a birth-death framework. PLoS Computional Biology 10, e1003771 (2014).

40. Finucane, H. K. et al. Partitioning heritability by functional annotation using genome-wide association summary statistics. Nature Genetics 47, 1228–1235 (2015).

41. Yang, Z. Maximum likelihood phylogenetic estimation from DNA sequences with variable rates over sites: approximate methods. Journal of Molecular Evolution 39, 306–314 (1994).

42. Tavaré, S. Some probabilistic and statistical problems in the analysis of dna sequences. Lectures on Mathematics in the Life Sciences 17, 57–86 (1986).

43. Pounds, S. & Morris, S. W. Estimating the occurrence of false positives and false negatives in microarray studies by approximating and partitioning the empirical distribution of p-values. Bioinformatics 19, 1236–1242 (2003).

44. ENCODE Project Consortium. An integrated encyclopedia of DNA elements in the human genome. Nature 489, 57–74 (2012).

45. Navarro Gonzalez, J. et al. The UCSC Genome Browser database: 2021 update. Nucleic Acids Res 49, D1046–D1057 (2021).

46. White, R.J. Transcription by RNA polymerase III: more complex than we thought. Nature Reviews Genetics 12, 459–463 (2011).

47. Boyer, L. A. et al. Core transcriptional regulatory circuitry in human embryonic stem cells. Cell 122, 947–956 (2005).

48. Liang, J. et al. Nanog and Oct4 associate with unique transcriptional repression complexes in embryonic stem cells. Nature Cell Biology 10, 731–739 (2008).

49. Arbiza, L. et al. Genome-wide inference of natural selection on human transcription factor binding sites. Nature Genetics 45, 723–729 (2013).

50. Gronau, I., Arbiza, L., Mohammed, J. & Siepel, A. Inference of natural selection from interspersed genomic elements based on polymorphism and divergence. Molecular Biology and Evolution 30, 1159–1171 (2013).

51. Gulko, B. & Siepel, A. An evolutionary framework for measuring epigenomic information and estimating cell-type-specific fitness consequences. Nature Genetics 51, 335–342 (2019).

52. McDonald, J. H. & Kreitman, M. Adaptive protein evolution at the Adh locus in drosophila. Nature 351, 652–654 (1991).

53. Smith, N. G. C. & Eyre-Walker, A. Adaptive protein evolution in Drosophila. Nature 415, 1022–1024 (2002).

54. McLean, C. Y. et al. Human-specific loss of regulatory DNA and the evolution of human-specific traits. Nature 471, 216 (2011).

55. Yu, G., Wang, L.-G., Han, Y. & He, Q.-Y. clusterprofiler: an r package for comparing biological themes among gene clusters. Omics: a journal of integrative biology 16, 284– 287 (2012).

56. Yu, G. & He, Q.-Y. Reactomepa: an r/bioconductor package for reactome pathway analysis and visualization. Molecular BioSystems 12, 477–479 (2016).

57. Capra, J. A., Hubisz, M. J., Kostka, D., Pollard, K. S. & Siepel, A. A model-based analysis of GC-biased gene conversion in the human and chimpanzee genomes. PLOS Genetics 9, e1003684 (2013).

58. Bernard, G. et al. Mutations of POLR3A encoding a catalytic subunit of rna polymerase pol iii cause a recessive hypomyelinating leukodystrophy. The American Journal of Human Genetics 89, 415–423 (2011).

59. Saitsu, H. et al. Mutations in POLR3A and POLR3B encoding rna polymerase iii subunits cause an autosomal-recessive hypomyelinating leukoencephalopathy. The American Journal of Human Genetics 89, 644–651 (2011).

60. Borck, G. et al. BRF1 mutations alter RNA polymerase III-dependent transcription and cause neurodevelopmental anomalies. Genome Research 25, 155–66 (2015).

61. Chew, J.-L. et al. Reciprocal transcriptional regulation of Pou5f1 and Sox2 via the Oct4/Sox2 complex in embryonic stem cells. Molecular and Cellular Biology 25, 6031– 6046 (2005).

62. Loh, Y.-H. et al. The Oct4 and Nanog transcription network regulates pluripotency in mouse embryonic stem cells. Nature Genetics 38, 431–440 (2006).

63. Lee, M. T. et al. Nanog, Pou5f1 and SoxB1 activate zygotic gene expression during the maternal-to-zygotic transition. Nature 503, 360–364 (2013).

64. Sharov, A. A. et al. Identification of Pou5f1, Sox2, and Nanog downstream target genes with statistical confidence by applying a novel algorithm to time course microarray and genome-wide chromatin immunoprecipitation data. BMC Genomics 9 (2008).

65. Leichsenring, M., Maes, J., Mössner, R., Driever, W. & Onichtchouk, D. Pou5f1 transcription factor controls zygotic gene activation in vertebrates. Science 341, 1005–1009 (2013).

66. Wang, J. et al. A protein interaction network for pluripotency of embryonic stem cells. Nature 444, 364–368 (2006).

67. Rodda, D. J. et al. Transcriptional regulation of Nanog by OCT4 and SOX2. Journal of Biological Chemistry 280, 24731–24737 (2005).

68. Lai, C. S., Fisher, S. E., Hurst, J. A., Vargha-Khadem, F. & Monaco, A. P. A forkhead-domain gene is mutated in a severe speech and language disorder. Nature 413, 519–523 (2001).

69. Fisher, S. E. & Scharff, C. FOXP2 as a molecular window into speech and language. Trends in Genetics 25, 166–177 (2009).

70. Vernes, S. C. et al. Foxp2 regulates gene networks implicated in neurite outgrowth in the developing brain. PLOS Genetics 7, e1002145 (2011).

71. Zhang, J., Webb, D. M. & Podlaha, O. Accelerated protein evolution and origins of human-specific features: Foxp2 as an example. Genetics 162, 1825–1835 (2002).

72. Enard, W. et al. Molecular evolution of FOXP2, a gene involved in speech and language. Nature 418, 869–872 (2002).

73. Atkinson, E. G. et al. No evidence for recent selection at FOXP2 among diverse human populations. Cell 174, 1424–1435.e15 (2018).

74. Li, G., Wang, J., Rossiter, S. J., Jones, G. & Zhang, S. Accelerated FoxP2 evolution in echolocating bats. PLOS ONE 2, 1–10 (2007).

75. Cahill, J. A. et al. Positive selection in noncoding genomic regions of vocal learning birds is associated with genes implicated in vocal learning and speech functions in humans. Genome Research 31, 2035–2049 (2021).

76. Li, Z., Cogswell, M., Hixson, K., Brooks-Kayal, A. R. & Russek, S. J. Nuclear respiratory factor 1 (NRF-1) controls the activity dependent transcription of the GABA-a receptor beta 1 subunit gene in neurons. Frontiers in Molecular Neuroscience 11 (2018).

77. Biswas, M. & Chan, J. Y. Role of Nrf1 in antioxidant response element-mediated gene expression and beyond. Toxicology and Applied Pharmacology 244, 16–20 (2010).

78. Meyer, A. G. & Wilke, C. O. Integrating sequence variation and protein structure to identify sites under selection. Molecular Biology and Evolution 30, 36–44 (2013).

79. Meyer, A. G., Dawson, E. T. & Wilke, C. O. Cross-species comparison of site-specific evolutionary-rate variation in influenza haemagglutinin. Philosophical Transactions of the Royal Society B: Biological Sciences 368, 20120334 (2013).

80. Huang, Y.-F. Dissecting genomic determinants of positive selection with an evolution-guided regression model. Molecular Biology and Evolution 39, msab291 (2022).

81. Hubisz, M. J., Pollard, K. S. & Siepel, A. PHAST and RPHAST: phylogenetic analysis with space/time models. Briefings in Bioinformatics 12, 41–51 (2011).

82. Schliep, K. P. phangorn: phylogenetic analysis in R. Bioinformatics 27, 592–593 (2011).

83. Cheng, R. Non-Standard Parametric Statistical Inference (Oxford University Press, 2017).

