## Supplementary table 1-3 for "Transcription factor binding sites are frequently under accelerated evolution in primates"

Supplementary Table 1: Non-overlapping TFBS groups under accelerated evolution. $r_1$ and $r_2$ are the relative substitution rates of a TFBS group in the human lineage and in other primates. $P$-values are calculated from likelihood ratio test. The ratio $(r_1/r_2)$ indicates the fold of increase in substitution rate in the human lineage.

| Genomic elements | $r_1$ | $r_2$ | *P*-value | $r_1/r_2$ |
|---|---|---|---|---|
| Pol III binding | 2.30 | 1.41 | 0 | 1.63 |
| BDP1 | 1.34 | 1.12 | 1.41E-06 | 1.19 |
| POU5F1-NANOG binding | 0.91 | 0.83 | 0.002 | 1.10 |
| POU5F1 | 1.03 | 0.94 | 1.5E-05 | 1.09 |
| FOXP2 | 0.99 | 0.92 | 0 | 1.08 |
| NANOG | 0.90 | 0.85 | 0.004 | 1.05 |
| NRF1 | 1.16 | 1.11 | 0.03 | 1.05 |

Supplementary Table 2: Relative BICs for different foreground lineages.

| Model | Foreground lineage | Pol III binding | BDP1 | POU5F1-NANOG binding | POU5F1 | FOXP2 | NANOG | NRF1 |
|---|---|---|---|---|---|---|---|---|
| M1 | Human | 158.29 | 23.27 | 9.82 | 18.74 | 141.34 | 8.24 | 4.79 |
| M2 | Hominini | 530.38 | 35.65 | 24.47 | 36.96 | 199.91 | 16.67 | 30.24 |
| M3 | Homininae | 1060.78 | 40.06 | 24.11 | 23.77 | 211.00 | 10.36 | 33.54 |
| M4 | Hominidae | 1590.02 | 65.20 | 15.94 | 17.96 | 320.93 | 20.07 | 46.08 |
| M5 | Caterrhini | 1973.45 | 113.72 | 8.81 | 36.88 | 186.56 | 13.15 | 47.28 |
| M6 | Simiiformes | 1110.76 | 58.78 | 2.09 | 20.54 | 28.66 | 0.13 | 6.30 |
| M7 | Haplorhini | 0 | 0 | 0 | 0 | 0 | 0 | 0 |

Supplementary Table 3: Rates of adaptive substitutions estimated by the INSIGHT model. $D_p$ indicates the expected number of divergences per kilobase driven by positive selection in the human lineage. SE$[D_p]$ is the standard error of $D_p$. *P*-values are calculated from the Wald test.

| Genomic elements | $D_p$ | SE$[D_p]$ | *P*-value |
|---|---|---|---|
| HARs | 7.67 | 0.14 | $< 0.0001$ |
| Pol III binding | 7.28 | 0.59 | $< 0.0001$ |
| BDP1 | 0.60 | 0.35 | 0.40 |
| POU5F1-NANOG binding | 0 | 0 | |
| POU5F1 | 0 | 0 | |
| FOXP2 | 0 | 0 | |
| NANOG | 0 | 0 | |
| NRF1 | 0.27 | 0.26 | 0.15 |
